# Context-aware knowledge selection and reliable model recommendation with ACCORDION

**DOI:** 10.1101/2022.01.22.477231

**Authors:** Yasmine Ahmed, Cheryl A. Telmer, Gaoxiang Zhou, Natasa Miskov-Zivanov

**Affiliations:** Computer Science Department, Virginia Tech, Blacksburg, VA 24060; Department of Biological Sciences, Carnegie Mellon University, Pittsburgh, PA 15213; Electrical and Computer Engineering Department, University of Pittsburgh, Pittsburgh, PA 15213; Electrical and Computer Engineering Department, Bioengineering, Computational and Systems Biology, University of Pittsburgh, Pittsburgh, PA 15213

**Keywords:** Knowledge and data engineering tools and techniques, Model checking, Patterns, Graphs and networks, Data models, Data/Text mining, Bioinformatics databases, Clustering, Query formulation, Natural Language Processing, Molecular Biology, Knowledge modeling, Simulation

## Abstract

New discoveries and knowledge are summarized in thousands of published papers per year per scientific domain, making it incomprehensible for scientists to account for all available knowledge relevant for their studies. In this paper, we present ACCORDION (**ACC**elerating and **O**ptimizing model **R**ecommen**D**at**ION**s), a novel methodology and an expert system that retrieves and selects relevant knowledge from literature and databases to recommend models with correct structure and accurate behavior, enabling mechanistic explanations and predictions, and advancing understanding. ACCORDION introduces an approach that integrates knowledge retrieval, graph algorithms, clustering, simulation, and formal analysis. Here, we focus on biological systems, although the proposed methodology is applicable in other domains. We used ACCORDION in nine benchmark case studies and compared its performance with other previously published tools. We show that ACCORDION is: *comprehensive*, retrieving relevant knowledge from a range of literature sources; very *effective*, reducing the error of the initial baseline model by more than 80%, recommending models that closely recapitulate desired behavior, and outperforming previously published tools; *selective*, recommending only the most relevant, context-specific, and useful subset (15-20%) of candidate knowledge in literature; *diverse*, accounting for several distinct criteria to recommend more than one solution, thus enabling alternative explanations or intervention directions.

## 1 Introduction

DISCOVERIES, predictions, design of treatments and interventions in biological and many other systems require understanding of *system dynamics*. To retrieve useful information and create reliable models for studying system dynamics, modelers often survey published papers, search model and interaction databases (e.g., Reactome [1], STRING [2], KEGG [3], etc.), incorporate background and common-sense knowledge of domain experts, and interpret data and observations from wet-lab experiments. Several million new scientific papers are published every year, with thousands in individual scientific domains, making it incomprehensible for scientists to account for all available knowledge relevant to their studies. The time-consuming manual steps make the creation of models a slow, laborious and error-prone process. On the other hand, machine learning and bioinformatics advances have enabled automated inference of network models from data. Although very proficient in identifying associations and correlations between system components, these methods still struggle if tasked with finding directionality of influences and causation [4], which are necessary in order to study system dynamics, the state changes in system and its components over time. Inferring large causal models from data requires significant time and computational resources, it is strongly dependent on the quality of the data, and on the applied statistics and machine learning methods [5]. The rapid growth of the amount of biological data in the public domain also aggravates the issues with data inconsistency and fragmentation [6]. Therefore, to efficiently create interpretable dynamic models, it is necessary to develop novel methods that combine (*i*) automated retrieval and selection of new, reliable, and useful information about component influences and causality, with (*ii*) automated recommendation of how to incorporate this information into models. Besides the significant speedup over slow manual steps, this would also expand the current capabilities for retrieval and processing of textual data and information about influences and causality. All of the above would in turn result in a *consistent, comprehensive, robust*, and *curated* process for creating dynamic models.

A recent review in [7] discusses methods that automate extension and recommendation of dynamic models by identifying and selecting relevant information among large sets of causal relationships, usually retrieved from literature [8][9][10][11]. While all methods reviewed in [7] succeed to some extent in addressing the above described challenges, each one of them still has drawbacks. They are either not scalable for large amounts of available information [8][9], non-deterministic (provide different solutions when run multiple times) [9], or attempt to create dynamic models based mainly on the static graph structure, not accounting for the dynamic behavior [10][11].

In this work, we propose ACCORDION (**ACC**elerating and **O**ptimizing model **R**ecommen**D**at**ION**s), a tool that identifies useful and relevant information from published literature and recommends model modifications that lead to closely recapitulating desired system behavior, all in a *fully automated manner*. Thus, compared to the work in [10], [11], ACCORDION also considers the dynamic behavior, and in contrast to [8][9], it focuses on identifying clusters of strongly connected elements in the newly extracted information that can have a measurable impact on the dynamic behavior of the model. ACCORDION is versatile, it can be used to extend any model that has a directed graph as an underlying structure (with the system components as nodes and the influences between components as directed edges), and update functions for elements, allowing studies of system dynamics. These models are often referred to as *executable models*. To demonstrate the efficiency and utility of the tool, we have selected nine different case studies using models of three systems, namely, the T cell differentiation model [12], the T cell large granular lymphocyte model [13], and the pancreatic cancer cell model [14], and seven machine reading outputs with varying features.

We show that ACCORDION can automatically, without human intervention, recommend new models that significantly reduce baseline model error and recapitulate known or desired system behavior. The contributions of the work presented here include:

1. *Recommendation* of executable dynamic models of cell signaling that satisfy known or desired system properties.
2. *Integration* of information retrieval, graph-based methods, and dynamic system analysis.
3. *“In-design” validation* of dynamic models, i.e., during their creation (instead of typical “post-design” approach, i.e., after models are created).
4. Rapid *exploration of redundancies* and the *discovery of alternative pathways* of regulation.
5. Execution of thousands of *in silico experiments* in at most a few hours, which would take days, or months, or would be impractical to conduct in vivo or in vitro.
6. *Open access ACCORDION tool*, that includes novel approaches and methods ((i)-(v) above), available on GitHub, with detailed documentation.

## 2 METHODS

Here, we first describe inputs to ACCORDION, followed by the description of the novel methodology within ACCORDION for processing these inputs to generate the output. Input and output examples and the flow chart of the entire approach are provided in Figure 1.

**Figure 1.**
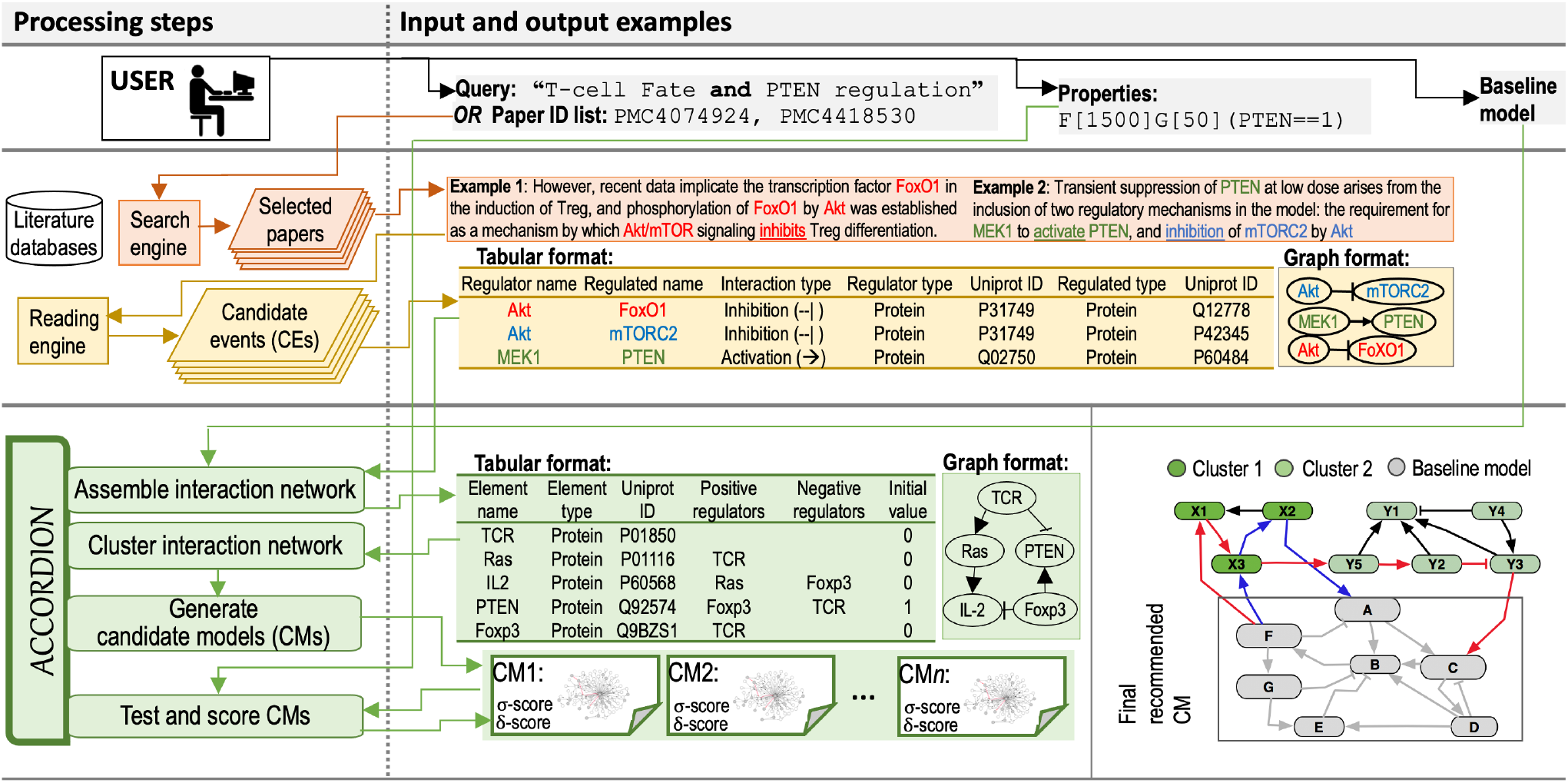
**Processing steps** column: The diagram of the flow and steps for information retrieval and processing, and model recommendation, including a user (**Top**), components of information retrieval from databases (**Middle**), and ACCORDION components (**Bottom**). **Input and output examples** column: (**Top**) Example query used to select relevant papers and example property in BLTL format. (**Middle**) Two example sentences with highlighted entities and events extracted by machine readers. Tabular outputs from REACH engine with **Example 1** and **Example 2** sentences as input. Graphical representation of REACH outputs. (**Bottom**) Tabular representation of several elements and their influence sets (positive and negative regulators) in Bio-RECIPES format [43] and the graphical representation of elements and influence sets. A toy example graph (*G*^*new*^) consisting of a baseline model and connected extension clusters: grey nodes belong to the baseline model, light and dark green nodes belong to the CE set obtained from machine reading, blue edges highlight a return path within one cluster, and red edges show a return path connecting two clusters. The multi-cluster path starts at Baseline model, continues through Cluster 1, then through Cluster 2, and ends in Baseline model.

### 2.1 Network and model inputs

#### Baseline model

One of the inputs to ACCORDION is a baseline model (Figure 1, top row, and gray nodes and edges in the bottom row), setting the context for other inputs and for the analysis. The baseline model can be created manually, with expert input, inferred automatically from data, or adopted from models published in literature [12][13][14][15] and in model databases [1][3][16]. To allow for studying dynamics, ACCORDION works with models that have *directed* graph structure, *G*(*V, E*), including both cyclic and acyclic graphs, where each node *v* ∈ *V* corresponds to one model element, representing a protein, gene, chemical, or a biological process, and each directed edge *e*(*v*_*i*_, *v*_*j*_) ∈ *E* indicates that element *v*_*j*_ is regulated or influenced, directly or indirectly, by element *v*_*i*_.

We refer to the set of regulators of an element as its *influence set*, distinguishing between positive and negative regulators. ACCORDION assigns to each element *v* a discrete variable *x*, which can be used to represent the element’s state, such as the level of its activity or amount. Each model element may have a state transition function, referred to as an *element update rule*, which defines its state changes given the states of its regulators, thus enabling the study of system dynamics. While the types of elements and their update rules are not constrained by the main methods implemented within ACCORDION (see Sections 2.2-2.3), they are largely affected by the information that is available in new events (see the following subsection) and in the baseline model. Most often, the events described in literature are qualitative, for example, only two element states (e.g., inactive/active, absent/present) may be distinguished or relevant, or only two or three levels of concentration may be considered (e.g., low/high or low/medium/high). Causal and Boolean types of regulations and update rules are most suitable in such cases, and ACCORDION is also compatible with such qualitative information. The details of model representation and formats accepted by ACCORDION are provided in the tool documentation [17][18].

#### Candidate event set

Another input to ACCORDION is a set of candidate events (CEs), which can be collected from different sources and created manually or automatically.

An example automated pipeline that can be used to create a CE set includes machine readers and INDRA (Integrated Network and Dynamical Reasoning Assembler) database of interactions extracted from literature. Machine reading of published literature with engines such as REACH [19] and TRIPS [20] can output large event sets, and therefore, allow for a high throughput processing of available information. INDRA [21] is an automated model assembly tool designed for biomolecular signaling pathway models and generalized to other domains such as disease models. INDRA relies on collecting and scoring new information extracted either from the textual evidence by machine readers or from structured pathway databases such as SIGNOR [22]. To select the most valuable and highquality statements, INDRA computes an overall belief score for each statement, defined as the joint probability of correctness implied by the evidence.

The set of relevant papers can be selected either using search tools such as Google or PubMed [23] or by providing key search terms to reading engines, which then directly use Medline search tools (e.g., PubMed [23], Ovid [24]) to find most relevant papers. Examples of queries, sentences processed by machine readers, and events in the machine reading output are shown in Figure 1. Each event in the CE set has a direction (source and target of interaction) and sign (positive or negative regulation).

### 2.2 Influence network recommendation

#### G^new^ creation and return path definition

The CE set can be represented as a set of directed edges, *E*^*CE*^, where the source and target nodes of these edges (events) form set *V*^*CE*^. From the baseline model (BM) graph *G*^*BM*^(*V*^*BM*^, *E*^*BM*^), and the CE set, ACCORDION creates a new graph *G*^*new*^(*V*^*new*^, *E*^*new*^), where *V*^*new*^ = *V*^*BM*^ ∪ *V*^*CE*^, and *E*^*new*^ = *E*^*BM*^ ∪ *E*^*CE*^. The edges *e*(*v*_*s*_, *v*_*t*_) in *E*^*CE*^, where *v*_*s*_ is the source node and *v*_*t*_ is the target node, can be classified into three categories: (*i*) both source node *v*_*s*_ and target node *v*_*t*_ are found in the baseline model: {*v*_*s*_, *v*_*t*_}*∈ V*^*BM*^; *(ii*) either the source node or the target node is found in the baseline model: (*v*_*s*_*∈ V*^*BM*^ and *v*_*t*_*∉ V*^*BM*^) or (*v*_*s*_*∉ V*^*BM*^ and *v*_*t*_*∈ V*^*BM*^); (*iii*) neither the source node nor the target node is found in the baseline model: {*v*_*s*_, *v*_*t*_}*∉ V*^*BM*^.

Adding the entire set of CEs to the baseline model all at once usually does not result in a useful and accurate model due to a very large ratio |*E*^*CE*^|/|*E*^*BM*^ |. Alternatively, we can add one interaction at a time and test each model version, which is time consuming, or even impractical, given that the number of models increases exponentially with the size of the CE set. Moreover, adding individual interactions does not have an effect on the model when an interaction is not connected with the model (case (*iii*) above). It proves most useful to add paths of connected interactions, which are at the same time connected to the baseline model in their first and last nodes. Therefore, our approach for finding the most useful subset of the CE set includes finding connected interactions, that is, a set of edges in the graph *G*^*new*^ that form a return path. We define a path of *k* connected edges as *e*^*path*^(*v*_*s*1_, *v*_*tk*_) = (*e*_*i*1_(*v*_*s*1_, *v*_*t*1_), *e*_*i*2_(*v*_*s*1,_ *v*_*t*1_), *e*_*i*3_(*v*_*s*3_, *v*_*t*3_), …., *e*_*ik*_(*v*_*sk*_, *v*_*tk*−1_, *v*_*tk*_)), and we will refer to *e*^*path*^(*v*_*s*1_, *v*_*tk*_) as a return path, when {*v*_*s*1_, *v*_*tk*_}*∈ V*^*BM*^ (Figure 1). ACCORDION searches for such return paths after clustering graph *G*^*new*^, which is discussed in the following.

#### G^new^ clustering

To find clusters in *G*^*new*^, we apply Markov Clustering algorithm (MCL) [25], an unsupervised graph clustering algorithm, commonly used in bioinformatics (e.g., clustering of protein-protein interaction networks [26][27]). A number of previous studies have demonstrated that the MCL algorithm outperforms other clustering techniques [26][28][29][30][31][32]. The MCL algorithm has been proven to converge with undirected graphs [25], and since in this early step we are interested in clustering a graph given its connectivity only, the information about adjacency without directionality is sufficient for this step. The directionality will be included in later steps when exploring dynamic behavior. Therefore, ACCORDION provides to the MCL algorithm the information about node adjacency in *G*^*new*^. Furthermore, since graph *G*^*new*^ can be either acyclic or cyclic, our work demonstrates a novel application of the MCL algorithm beyond its previous use on acyclic graphs only [33].

MCL simulates random walks on an underlying interaction network (in our case, graph *G*^*new*^), by alternating two operations, expansion and inflation. First, self-loops are added to *G*^*new*^, and the updated graph is represented as an adjacency matrix M, which is therefore symmetric, mapping nodes in *G*^*new*^ to both row and column headers in M. The entries in matrix M are assigned value 1 when an edge between their column and row nodes exists in *G*^*new*^ or when an entry is on the main diagonal of M (i.e., same column and row node), and value 0 otherwise. Next, matrix M is used by the MCL algorithm as an initial version of a stochastic Markov matrix M′ [34], where each entry represents the probability of a transition from the column node to the row node. Since *G*^*new*^ is not a weighted graph, all transitions are assumed to be equally likely, and the matrix M′ is normalized such that the sum of entries in each column equals 1.

The probability of a random walk of length *q* between any two nodes can be calculated by raising the matrix M′ to the exponent *q*, a process called “expansion”. As the number of paths is likely larger between nodes within the same cluster than between nodes across different clusters, the transition probabilities between nodes in the same cluster will typically be higher in a newly obtained expanded matrix. MCL further amplifies this effect by computing entry-wise exponents of the expanded matrix, a process called “inflation” [25], which raises each element of the matrix to the power *r*. Clusters are determined by alternating expansion and inflation, until the graph is partitioned into subsets such that there are no paths between these subsets. The final number of generated clusters, *C*_1_, …, *C*_*n*_, depends on the selected inflation parameter *r* [25].

As discussed above, ACCORDION clusters the entire *G*^*new*^ in order to account for the connectivity between new elements in CE and the baseline model, and thus, it likely assigns parts of the baseline model to different clusters. We will refer to the CE (BM) part of a generated cluster *l* as 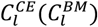 and to the nodes and edges in these cluster subsets as 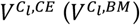 and 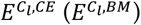, respectively.

#### Assembly of candidate influence networks

From the generated clusters and the baseline model, ACCORDION assembles multiple *candidate models* (*CMs*) as follows. ACCORDION can add clusters one at a time, or in groups. The more clusters or cluster groups are generated, the number of possible cluster combinations grows, and consequently, ACCORDION needs to assemble and test more models. In addition to that, in most cases |*V*^*BM*^ | < |*V*^*CE*^| and |*E*^*BM*^ | < |*E*^*CE*^|, and thus, the number of new nodes and edges in a cluster tends to be relatively large compared to the size of the baseline model (we will show related examples for our case studies later in Section 3).

Adding a large number of new nodes and edges to the baseline model at once can significantly change the structure, and consequently, the behavior of the model. Therefore, the default approach in ACCORDION is to evaluate only individual clusters generated as described in previous sub-section, as well as merged clusters *C*_*i,j*_, created by combining pairs of clusters *C*_*i*_ and *C*_*j*_ (*i, j* = 1, … *n, i* ≠ *j*). ACCORDION determines for each individual and merged cluster whether it forms a return path with the baseline model, and for each such cluster, ACCORDION creates a candidate model by adding the entire baseline model to the cluster. In other words, the number of created candidate models is equal the number of clusters (individual and merged) that form a return path with the baseline model.

As defined above, the clusters formed from the *G*^*new*^ graph can contain nodes and edges of the baseline model. Therefore, for those clusters (individual or merged) that were used to create candidate models, ACCORDION computes the *node overlap* (*NO*) value [11], as a ratio of those nodes in a cluster C_4_ that are present in the baseline model, 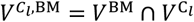 and the total number of nodes within the cluster 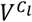:

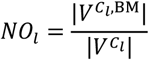

### 2.3 Executable model recommendation

In Section 2.2, we discussed the steps to form *G*^*new*^, focusing on its *static* structure. Here, we describe creation of new update functions for elements in *G*^*new*^, and how an additional input to ACCORDION is used to evaluate *dynamic* behavior of candidate models.

#### Updating element update rules

When adding new elements and influences to baseline models, ACCORDION uses the information provided in its inputs to update existing or create new element update rules. This information includes element update rules in the baseline model and the sign of influences (positive or negative) in the CE set. Whenever a new element *v* ∈ *V*^*CE*^\*V*^*BM*^ with non-empty influence set is added to the baseline model, ACCORDION generates a new update rule for *v*. While the algorithms within ACCORDION are not dependent on the type of state update rules used for elements of a baseline model (as discussed in Section 2.1), the selection of CMs will depend on the granularity of the information provided at the input. The event information available in the CE set is often qualitative, for example, “*A* positively regulates *B*”. Furthermore, if an update rule for element *B* in the baseline model already includes two positive regulators *C* and *D*, i.e., *x*_*B*_ = *f*(*x*_*C*_, *x*_*D*_), then the new event from the CE set (“*A* positively regulates *B*”) can be added to the update rule for *B* in two ways: using “OR” operation, *x*_*B*_ = *f*(*x*_*C*_, *x*_*D*_) OR *x*_*A*_, or using “AND” operation, *x*_*B*_ = *f*(*x*_*C*_, *x*_*D*_) AND *x*_*A*_ (following the definition from Section 2.1, *x*:, *x*_*B*_, *x*_*C*_, *x*_*D*_ are variables representing level or amount or activity of elements *A, B, C, D*, respectively). Additionally, when elements have more than two discrete levels, ACCORDION can, for example, use max and min operators to determine the maximum or minimum influence from a given set of regulators.

#### Model evaluation

The third input to ACCORDION includes a set of properties 𝓉 ∈ 𝒯, which define the dynamic behavior that the models recommended by ACCORDION should satisfy. We refer to this behavior as “desired behavior” and, dependent on the goals of a study, this can be actual, observed, measured, or expected behavior of the modeled system. To select the CM that allows for most closely reproducing the experimentally observed or desired behavior, and given the randomness in time and order of events in modeled systems, ACCORDION uses a combination of stochastic simulation and statistical model checking.

The DiSH (discrete stochastic heterogeneous) simulator [35][36] is used to obtain element trajectories, i.e., a sequence of element state values in time, for the baseline model and the CMs. DiSH is a stochastic simulator that can simulate models at different levels of abstraction, information resolution, and uncertainty. This range of simulation schemes is especially valuable when working with diverse information sources and inputs, such as the ones used by ACCORDION. Each simulation run starts with a specified initial model state, where initial values are assigned to all model elements to represent a particular system state (e.g., naïve or not differentiated cell, healthy cell, cancer cell). The initial values for the baseline model elements (nodes in *V*^*BM*^) are usually already known, however, the newly added elements (nodes in *V*^*CE*^) need to be assigned initial values as well. For the purpose of presenting ACCORDION here, we assume that, when no initial values are provided for new elements, all elements in the same cluster start at a similar level. As our future step, we will expand ACCORDION with several methods for inferring and assigning initial values, if data is available.

ACCORDION runs a statistical model checker [37][38] to verify whether the CMs satisfy a set of desired system properties. The model checker reads properties formally written using Bounded Linear Temporal Logic (BLTL) [39][38] and, for a given executable model ℳ and a property 𝓉, it outputs a *property probability estimate*, 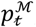, that ℳ satisfies 𝓉, under predefined error interval for the estimate. For instance, we can test whether at any point within the first *s*_1_ time steps, model element *v*_*i*_ (i.e., its state variable *x*_*i*_) reaches value X_1_ and element *v*_*j*_ (i.e., its state variable *x*_*j*_) reaches value X_1_, and they both keep those values for at least *s*_1_ time steps. We write this property formally as 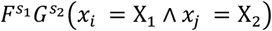, where 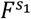 stands for “any time in the future *s*_1_ steps”, and 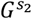 stands for “globally for *s*_1_ steps”. An example of a property and its expected value is shown in Figure 1. To avoid the search for all possible state trajectories through the non-deterministic state transition graph, the statistical model checker calls the simulator to generate element trajectories for a defined number of steps and performs statistical hypothesis testing on those trajectories with respect to a given property [40][8].

#### Model scoring and recommendation

Generated CMs can be scored in different ways, depending on the goals of the study. Once all created CMs are evaluated on how well they satisfy each given property, ACCORDION can find models that satisfy a particular property 𝓉_*j*_ ∈ 𝒯 with high probability. To provide the recommendation of top CMs that are closest to expected probability values for properties, we use several metrics defined as follows.

*Definition 1*: The *goal property probability* for a property 𝓉, denoted as P_𝓉_, indicates either the estimated likelihood or expected likelihood (e.g., after an intervention) for the real system to satisfy property 𝓉.

We note here that, due to randomness of biological systems, P_𝓉_ is not always 0 or 1, and instead can take any value from the interval [0,1].

*Definition 2*: For a given model ℳ and property 𝓉, the *model property error*, 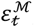, is the absolute difference between the probability value 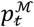 estimated by model checking for model ℳ and property 𝓉, and the goal property probability 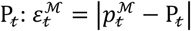.

*Definition 3*: For a given model ℳ, the *average model error*, 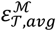, is computed as a mean of model property errors 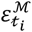 across all tested properties 𝓉_*i*_ ∈ 𝒯.

*Definition 4*: For a given model ℳ and a set of properties 𝒯, we define *α-score* as 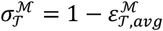.

It can be concluded from Definition 4 that the larger the *α*score for a model is, the closer the model is to satisfying all desired properties.

*Definition 5*: For a given model ℳ, a set of properties 𝒯, and δ ∈ [0,1], we define *δ-score*, denoted as 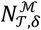, as the percent of properties in 𝓉_*i*_ ∈ 𝒯 for which it holds that 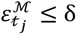.

The parameter δ indicates how close the 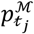 value needs to be to the goal probability 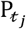 for the property to be considered satisfied. ACCORDION users can select the value for δ based on their modeling goals.

## 3 Results

### 3.1 Benchmarks

In the absence of standardized benchmarks to evaluate ACCORDION, we created nine case studies. These benchmarks and all related files will be open access and available with ACCORDION release [17][18]. In the Appendix, we provide an overview of the biological background for all studied systems, the details of creating the baseline model, and the steps of selecting literature and creating CE set for each conducted case study. In Figure 2 (a) and (b), we list the main characteristics of these nine cases, with models of three biological systems and different sets of CEs for each system. The three models include control circuitry of naïve T cell differentiation (T cell) [12], T cell large granular lymphocyte (T-LGL) leukemia model [13], and pancreatic cancer cell model (PCC) [14]. The studies vary in the size and graph features of baseline models (“BM creation” columns) and the CE sets (“CE set creation” columns), and are named Tcell CE^FA^, Tcell CE^SA^, Tcell CE^SM^, T-LGL Q^Sm^, T-LGL Q^Med^, T-LGL Q^Det^, PCC BM^Au^, PCC BM^Ap^, and PCC BM^Pr^ (see Appendix for details). The size of baseline models varies from several tens to several hundreds of nodes or edges, and the number of interactions in the CE sets varies from half the number of interactions in the baseline model to six times larger (“BM and CE set relationship” columns, Figure 2(a)). In Figure 2(c), we illustrate the overlap and differences between the CE sets in the T cell case studies, to highlight the variability across CE sets that can be obtained in the context of the same baseline model.

**Figure 2.**
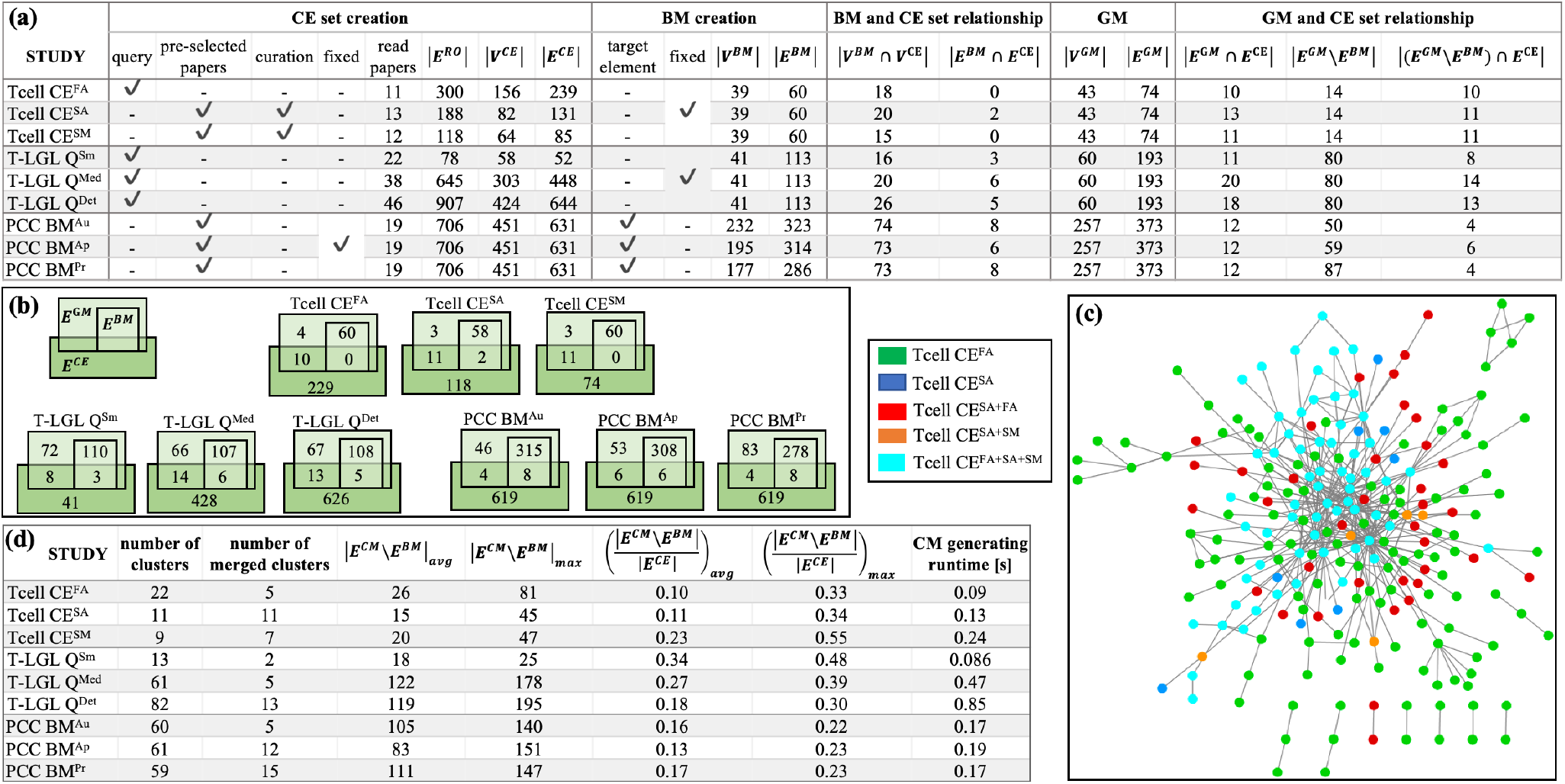
**(a)** Benchmark characterization: CE set creation procedure – using a query or a preselected set of papers, or manually curating the paper selection, and using a fixed or different CE set across all models for the same biological system; *E*^AB^ – number of events in the reading output; baseline model (BM) creation – fixed or different across all three studies for the same biological system; the intersection between BM and CE node and edge sets; the golden model (GM) *V*^*GM*^ and *E*^*GM*^ sizes; the relationship between each GM and the corresponding CE set (the number of common edges, the number of edges that are in GM but not in BM, the number of edges that are in GM but not in BM and are found in the CE). (**b**) Venn diagrams illustrating the overlap between three sets, *E*^*CE*^, *E*^*BM*^ and *E*^*GM*^ for the nine case studies. **(c)** Overlap and differences between the three CE sets for the T cell studies (legend on the left side of the figure). **(d)** Several characteristics of CMs created by ACCORDION for the nine case studies, including the runtime of ACCORDION for obtaining these CMs.

We also list in Table S1 in the Appendix the sets of properties that the real system satisfies, or should satisfy, which are not fully satisfied by baseline models and are used to guide new model assembly for each case study. The properties in Table S1 are provided in both natural language descriptions and in machine readable BLTL format, together with their goal probability values 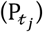. For each system, besides a baseline model, we also found a golden model in literature ([41] for the T cell model, [13] for the T-LGL model, and [14] for the PCC model). The details of each golden model are provided in the Appendix and as part of benchmark descriptions in the tool documentation [17][18]. Figure 2(a) includes the characteristics of golden models (columns “GM” and “GM and CE set relationship”).

With the nine case studies, we evaluate ACCORDION’s performance and demonstrate different use cases by: (*i*) varying the size and contents of the baseline model and the CE set (all nine case studies); (*ii*) varying the quality of the CE set (Tcell case studies); (*iii*) varying the level of detail in user selection of literature (Tcell CE^FA^ study and all three TLGL case studies); (*iv*) reconstruction of previously published model (all nine case studies).

We summarize in the table in Figure 2(d) the overall graph characteristics of the CMs obtained by ACCORDION for these nine benchmarks.

### 3.2 Recommending new models with desired behavior

The results listed in Figure 3 emphasize the importance of using ACCORDION when recommending a new or extended model. We show in Figure 3 the node overlap values (*NO*), and normalized values for *σ*-score, *δ*-score, and joint property probability estimate across all properties in 𝒯 (definitions in Section 2.3 and Appendix). These charts demonstrate that using only one of the metrics may be misleading since the “best” recommended CM can be different across these four metrics (differences highlighted in yellow in Figure 3). Furthermore, the results show that in some case studies ACCORDION recommends multiple CMs even when using the same metric (e.g., T-LGL Q^Med^ and Q^Det^ for all the metrics). Additionally, we show in the form of heatmaps, the individual property probability estimates 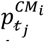 that ACCORDION computed in all nine case studies, for each tested property, for the CMs that formed return paths with baseline models. This detailed information can be especially useful if users decide to manually inspect and further modify CMs recommended by ACCORDION.

**Figure 3.**
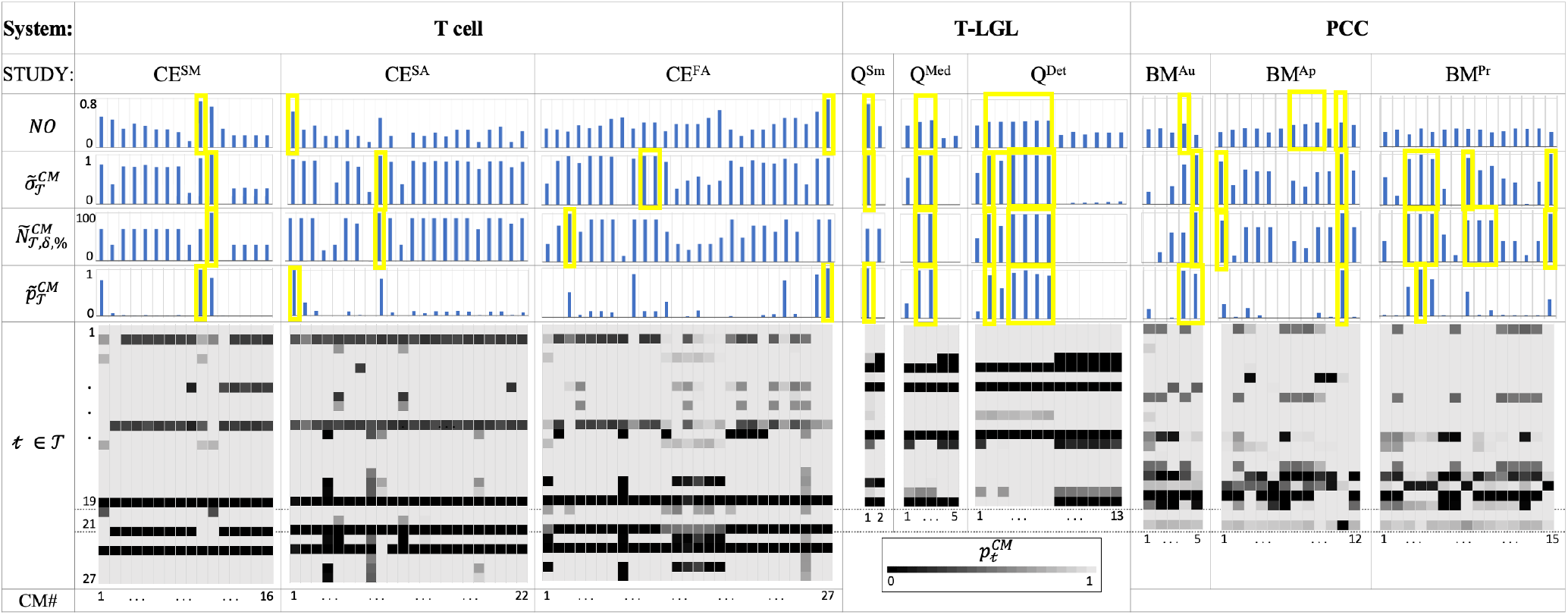
The Tcell, T-LGL and PCC use case results. For each CM (columns in heatmaps and bar charts): *NO* values; normalized 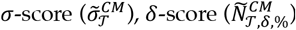, and joint property probability estimate 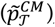; and heatmaps of individual property probability estimates for 27, 19 and 21 properties of the Tcell, T-LGL and PCC use cases, respectively. Results are shown for 16 (CE^SM^), 22 (CE^SA^) and 27 (CE^FA^) CMs for the Tcell studies, 2 (Q^Sm^), 5 (Q^Med^) and 13 (Q^Det^) CMs for the T-LGL studies, and 5 (BM^Au^), 12 (BM^Ap^) and 15 (BM^Pr^) CMs for the PCC studies. Normalized versions for the metrics (equations in Appendix) are used to clearly distinguish the “best” CM per each metric. Bars highlighted in yellow show that different metrics can recommend different CMs.

In Figure 4(a), we show the δ-score, 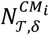, values for the top CMs recommended by ACCORDION in each of the nine studies. Additionally, for the parameter δ, which indicates the allowed difference between the computed CM property probability value and the goal probability, we explored a range of values (0, 0.1, 0.2, 0.3, 0.4, 0.5). To highlight the improvements in CMs when compared to the original baseline model, we show all results next to their corresponding baseline model values. ACCORDION achieved δ-score of 95% when δ=0.3 (all but one property is satisfied for this value of δ). Furthermore, increasing δ improves the model score, however, we observed that 0.2 or 0.3 value for δ is optimal to obtain useful models with high score. Overall, ACCORDION *automatically selected a small fraction (∼20%) of all interactions in the CE set, sufficient to decrease model error by up to 83%* (Figure 4(b)). Ideally, we would like the baseline model error to be reduced to 0, however, our case studies were designed to mimic realistic scenarios, where the recommended model error can be affected by external factors, as discussed in the following.

**Figure 4.**
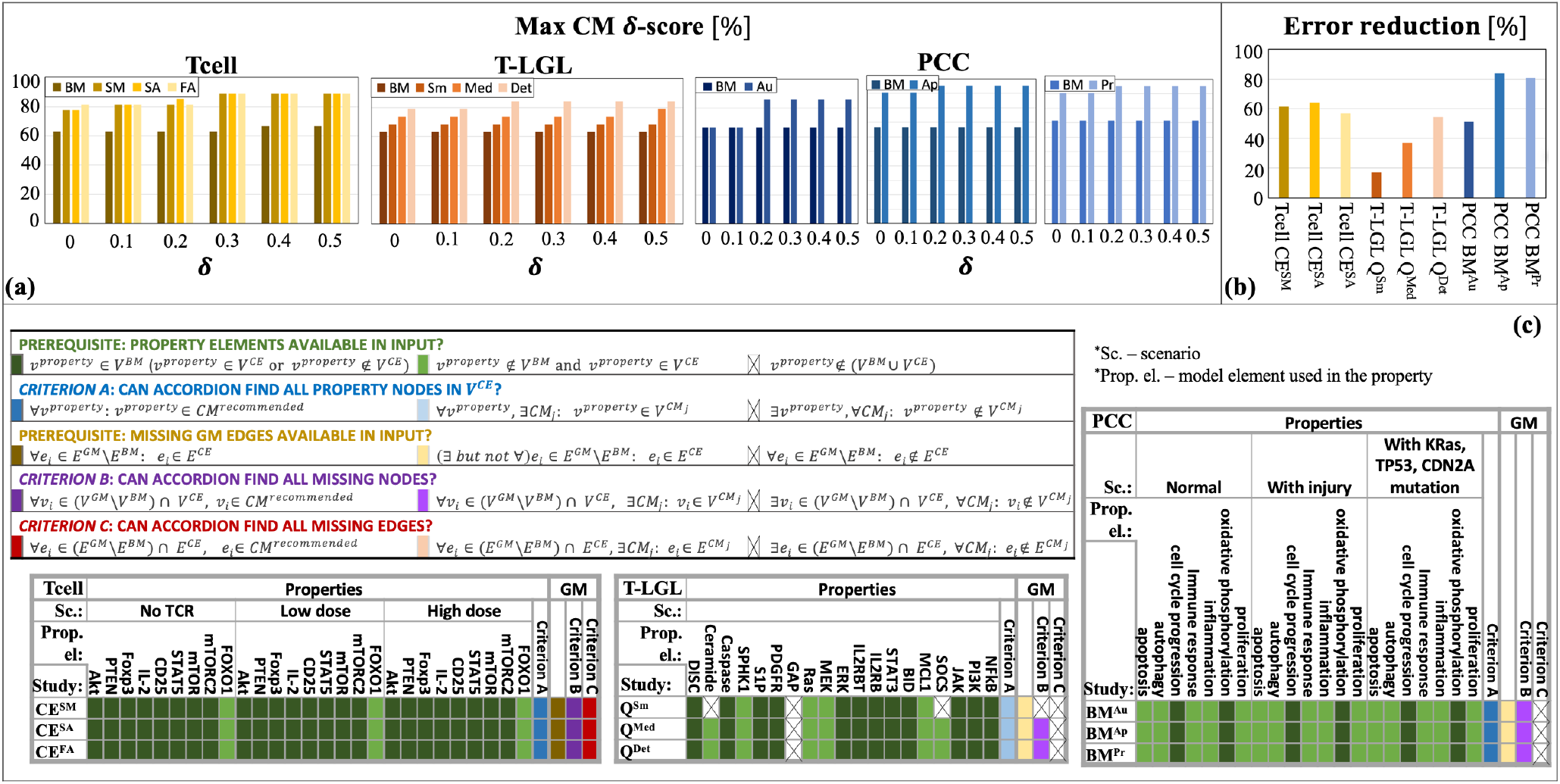
ACCORDION evaluation on nine case studies, Tcell studies SM, SA, FA, T-LGL studies Sm, Med, Det, and PCC studies Au, Ap, Pr: (a) maximum CM 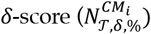obtained in each case study, compared with the baseline model (BM) *δ*-score; the results are compared for different values of *δ* (0, 0.1, 0.2, 0.3, 0.4, 0.5); (b) error reduction ACCORDION achieves in each case study; (c) definitions of two prerequisites for using properties (green) or reconstructing golden models (orange), and three criteria for evaluating ACCORDION’s outcomes (blue, purple, and red), including possible cases (shades of each color) (*v*^*property*^ is the element included in the property, property details listed in Appendix; *CM*^*recommended*^ is the top recommended model); tables show whether the prerequisites or criteria are satisfied for all nine case studies.

Several CE sets did not fulfill the necessary requirement for properties to be used: all elements that are listed in properties (Table S1, Appendix) need to be present in at least one of the sets *V*^*BM*^ and *V*^*CE*^. As shown in Figure 4(c) (“Properties” columns, green), in six out of nine studies, these elements are either already in the baseline model or in the CE set. However, in all three T-LGL studies, element GAP is not found in either of the two sets, *V*^*BM*^ and *V*^*CE*^, and in the T-LGL Q^Sm^ case additional two elements, Ceramide and SOCS, are not present. These element omissions occur in ACCORDION’s input and are due to machine reading not finding those elements in selected papers. While the properties that correspond to such omitted elements are not suitable for evaluating ACCORDION, we included them in our results to demonstrate realistic cases with imperfect CE sets. As part of our future work on ACCORDION, we plan to include pre-processing methods to automatically exclude such tests before clustering the CE set, or to inform the user at the beginning that property elements are not found in the input.

While it is not reasonable to expect from ACCORDION to find an element that does not exist in its input, it should be able to recover property elements that are not present in *V*^*BM*^ but are found in input *V*^*CE*^. As Figure 4(c) (“criterion A”, blue) highlights, ACCORDION *does indeed recover all property elements missing from a baseline model in at least one of the recommended CMs*.

Finally, when ACCORDION recovers all necessary property elements, most often the reason for non-zero model property errors 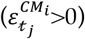 is in element update rules. For instance, in the Tcell cases, for the best recommended model per case, ACCORDION was able to recover FOXO1 which was not in *V*^*BM*^ but was in *V*^*CE*^. Moreover, ACCORDION recovered the update function of FOXO1 in all three cases and therefore, the properties that correspond to the dynamic behavior of FOXO1 (𝓉_9_, 𝓉_18_ and 𝓉_27_) under three different scenarios were all satisfied by most CMs (heatmaps, Figure 3). However, in the case of update function for AKT, ACCORDION added a number of new AKT regulators to the baseline model which affected the dynamic behavior of AKT. There are two ways in which this could be overcome. First, one could either use other tools to filter or score individual interactions in CE set [21][42] *before* they are used by ACCORDION, which we are planning to incorporate as one of our future steps. Second, ACCORDION can be used to identify cases where human input is necessary, for example, cases where many element regulators appear in literature, but not all of which can be used to form regulatory rules.

### 3.3 Finding most relevant set of new interactions

To test how well ACCORDION performs under a range of different conditions, we created the use cases such that the relationship between the number of elements and interactions in baseline models (|*V*^*BM*^|, |*E*^*BM*^ |), and in their corresponding CE sets (|*V*^*CE*^|, |*E*^*CE*^|) varies, from the CE set being smaller than baseline model in the T-LGL Q^Sm^ case, to being up to six times larger than baseline model in other use cases (Figure 2). We also determined the size of the overlap, |*V*^*BM*^ ∩ *V*^*CE*^| (see Figure 2(a)), further highlighting that indeed the number of new elements that could be added to the model is much larger than the number of elements in the model.

Additionally, we created these nine case studies such that they have baseline models with varying level of network connectivity. The baseline model in the T cell studies (Case studies section, Appendix), is a previously published, thus functional, model, while the T-LGL and PCC baseline models were created by removing nodes and interactions from published models. Since by construction the clusters that ACCORDION generates are connected only to a part of the baseline model (Section 2.2), we used the node overlap metric *NO* to determine the relationship between the number of new nodes that are added to the baseline model and the part of the model these nodes are connected to. The *NO* numbers in Figure 3, together with the ratios 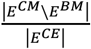 listed in Figure 2(d), show that ACCORDION is very selective, and it *only adds to the baseline model a subset of new interactions that are well connected with the baseline model*.

We investigated the percentage of these interactions selected from the entire CE set that were included in the top recommended CM (Figure 2(d)). For the Tcell cases, ACCORDION recommended on average 14% of the interactions as candidates for model extension, whereas for T-LGL and PCC cases, ACCORDION identified on average 26% and 15% of such interactions, respectively. These numbers emphasize an important characteristic of ACCORDION: *while allowing for comprehensive overview of literature, it significantly reduces the number of selected interactions, such that, if human input is still necessary, the number of interactions to manually review is significantly smaller than the original CE set*. Interestingly, higher *NO* values seem to correlate well with larger reduction in model error for the Tcell and TLGL studies. However, in the PCC studies this correlation does not hold, where the CMs with a large number of new interactions compared to the size of the baseline model significantly decrease the baseline model error (∼80% reduction, as shown in (Figure 4(b)). This demonstrates another important outcome: *when the baseline model is complete and well tested a smaller number of extensions can help improve it (e*.*g. Tcell and T-LGL cases), while for baseline models that are incomplete (e*.*g*., *when the user starts only with a seed set of interactions and not a complete model), a larger number of interactions needs to be added to improve them (e*.*g*., *PCC case)*.

### 3.4 Identifying alternative networks

As described in Section 3.1, besides baseline models, we also used golden models in our case studies. The purpose of comparison with golden models is to (*i*) determine how closely ACCORDION can reproduce previously published models (“criterion B”, purple, and “criterion C”, red, in Figure 4(c)) and (*ii*) what other models, different from golden models and satisfying the same set of properties, ACCORDION is able to create.

In all three T cell case studies, ACCORDION adds all the interactions from the *E*^*GM*^\*E*^*BM*^ set to its top recommended CMs (columns “GM” in Figure 4(c), orange). For example, one of the merged clusters in the Tcell case, with *NO*=0.7, restored all the missing interactions that were removed from the golden model. In the T-LGL and PCC studies, ACCORDION adds 30% and 32% of missing golden model interactions to recommended CMs. Similar to the discussion in Section 3.2 about the presence of property elements in *V*^*BM*^ and *V*^*CE*^, only interactions that are present in the input CE set can be examined by ACCORDION (prerequisite in Figure 4(c), orange). To this end, we find that in all three Tcell studies all golden model interactions that are missing from the baseline model, i.e., interactions from the *E*^*GM*^\*E*^*BM*^ set, are present in CE sets. On the other hand, the CE sets in the T-LGL and PCC studies do not contain all the interactions from the *E*^*GM*^\*E*^*BM*^ sets, (Figure 4(c), columns “GM”, orange). There are two possible reasons for this, either papers that were selected using queries do not include those missing interactions or machine reading does not recognize these interactions in the papers. An important outcome from this exercise is that ACCORDION recommends new CMs, different from golden models, which have high *σ*-score and δ-score and contain new interactions that form return paths with the baseline model. Moreover, in the T-LGL studies, a significant portion of interactions (41%) was removed from the golden model to obtain the baseline model. In such cases, ACCORDION selected from the large CE sets many additional interactions that form stronger connections with the baseline model (as part of clusters with high *NO* values and return paths) than the ones that are in the golden model, while also being able to find CMs that have high *σ*-score and δscore. For instance, the regulators of AKT in the golden model are PIP3 and mTORC2, while the models recommended by ACCORDION also include regulations by TGFB, IFNgamma, CK2, CTLA4, SHIP1, all of which are suggested in literature. *This highlights another possible use of ACCORDION: for examining redundancies in signaling networks or discovering alternative pathways regulating the same target element*.

### 3.5 Assistance in query answering

We also explored the relationship between the design of queries and ACCORDION’s effectiveness, that is, whether the selection of search terms to mine literature affects the usefulness of extensions selected by ACCORDION. For the Tcell CE^FA^ case, we used a search query as an input to PubMed to identify the most relevant papers (Case studies section, Appendix). We investigated the influence of this query on the percentage of interaction in clusters used to create CMs with top scores. In Figure 2(d), we show the average and the maximum percentage of selected interactions, i.e., 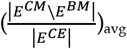 and 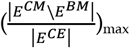, which are 10% and 33%, respectively. For the best recommended model of this particular case study, ACCORDION was able to recover all the missing elements that are in *V*^*GM*^ and not in *V*^*BM*^, namely, FOXO1, NEDD4, CK2 and MEK1. Furthermore, as can be seen in Figure 3, ACCORDION recapitulated the dynamic behavior of FOXO1, an element that was in the search query used to collect interactions for the CE set (Case studies section, Appendix), in all three scenarios (properties 𝓉_9_, 𝓉_18_and 𝓉_27_). However, the dynamic behavior of AKT (also in the search query), IL2 and STAT5 was not recovered in one out of three scenarios, (“High TCR” scenario in Table S1, Appendix, properties 𝓉_19_, 𝓉_21_and 𝓉_24_). This is due to erroneous interactions in the CE set extracted by machine readers. As mentioned previously, we plan to add a pre-processing of CE sets before using them with ACCORDION (e.g., using interaction filtering [42]).

For the T-LGL model study, we used three different queries (Case studies section, Appendix). The most elaborate query, in the T-LGL Q^Det^ case study, introduced more descriptive search terms, led to selecting more relevant papers, and consequently, extraction of relevant events and element regulators resulting in recommendation of a CM with high *σ*-score (0.76) and δ-score (0.75). Additionally, the update rules of most of the elements were retrieved except three elements, S1P, GAP and IL2RB. The properties that correspond to these three elements are properties 𝓉_5_, 𝓉_7_ and 𝓉_/1_. In contrast, for T-LGL Q^Sm^ and T-LGL Q^Med^ cases, less properties have been satisfied. For example, the baseline model error in property 𝓉_17_, related to the behavior of element JAK, is not corrected in the T-LGL Q^Sm^ case, while property 𝓉_19_, related to element NF*κ*B, is not corrected in both T-LGL Q^Sm^ and T-LGL Q^Med^ cases. This is mainly due to the key regulatory interactions for these elements not being extracted from literature, or due to the interactions that are recovered not forming proper update functions. Overall, by comparing the results for the three queries in the TLGL case studies, we have confirmed that a better query design leads to more useful and relevant information in the input CE sets.

### 3.6 Runtime and choice of inflation parameter

In Figure 2(d), we list the time that ACCORDION takes to generate clusters when run on a 3.3 GHz Intel Core i5 processor. The time required by ACCORDION to generate clusters increases with larger CE sets. For the PCC case studies, the runtime is same across studies since the same CE set has been used. However, for the T cell and T-LGL case studies, the CE sets have different sizes, and thus, result in different runtime. The runtime of the overall extension algorithm is proportional to the number of properties that we need to test against. In other words, if we have *N*_*L*_ clusters and *N*_*P*_ properties, the time required for the extension algorithm is at the order of *O*(*N*_*L*_ * *N*_*P*_). However, the runtime can be significantly reduced if testing for all properties and clusters is carried out in parallel, which is part of our immediate future work.

As we see above, the runtime is dependent on the number of clusters, which in turn is dependent on the cluster granularity and parameter values chosen for the MCL algorithm. The principal handle for changing cluster granularity is the inflation parameter *r*, described in Section 2.2.

An increase in *r* causes an increase in the cluster granularity. In [25], the authors determined a good interval to choose from (e.g., from 1.1 to 10.0), however, the range of suitable values also depends on the input graph. We explored the effect of *r* on finding the best set of clusters for each benchmark CE set. In Figure 5(a), we show our results for the Tcell case study and the different reading output sets, we found that *r*= 1.1 is too low, and *r* ≥ 6.0 is too high. We have therefore chosen value *r* = 4 for our studies and conducted the experiments discussed in previous sections using this value.

**Figure 5.**
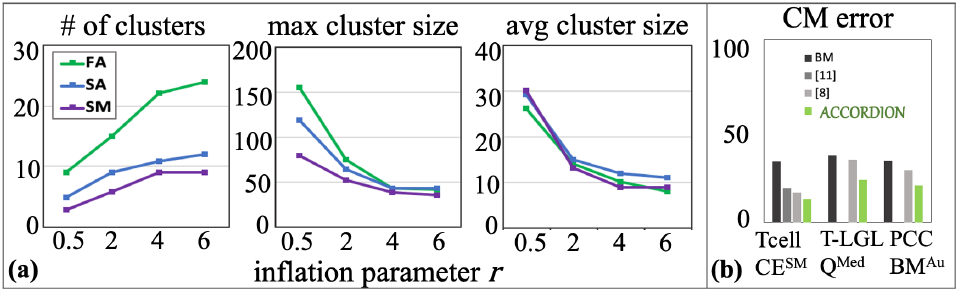
(a) Several cluster characteristics measured as functions of inflation parameter (*r*), for the Tcell CE^FA^, Tcell CE^SA^, and Tcell CE^SM^ cases (*r*_*1*_=0.5, *r*_*2*_=2, *r*_*3*_=4, *r4*=6). (b) The comparison between BM error and the top model (with minimum CM error, 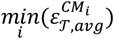, recommended by ACCORDION and other previously published methods in [8] and [11].

### 3.7 Comparison with other existing methods

We compare here the performance of ACCORDION with other model extension approaches. The method proposed in [8] iteratively expands a baseline model, by examining a large machine reading output in each iteration, and automatically selects a subset of interactions (influences) that can be directly connected with the baseline model. The work in [8] both expands the model network and tests the dynamics of the newly built model, by comparing it with a set of requirements or desired system states. The main drawback of the method in [8] is that it becomes impractical for large models due to adding new interactions in layers, based on their proximity to the existing model. On the other hand, the method proposed in [9] uses a genetic algorithm to select a set of extensions from machine reading output to create a new model with desired behavior. The two main disadvantages of this approach are issues with scalability and the non-determinism, as the solution may vary across multiple algorithm executions on the same inputs.

In [10][11], the authors proposed a tool and several metrics that rely on interaction occurrences and co-occurrences in published literature, and account for the connectivity of the newly added interactions to the existing models. While it selects new high-confidence interactions that are well supported by published literature and connected to the baseline model, this tool focuses on the static model network and does not consider its dynamic behavior.

We compared ACCORDION’s performance in terms of average model error of the top recommended model 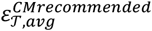 with two other previously published methods for model extension from [8] and [11] (the authors in [11] demonstrated that their methods outperform the methods from [9], thus, we chose to compare here only with the highest performing methods). Figure 5(b), shows that among all model extension methods, ACCORDION is able to find models with the lowest 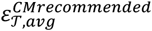. We applied the layered approach from [8] only on the Tcell case study, since it has been shown to mainly work on smaller models, and we applied the approach from [11] on all three baseline models. The method in [11] relies only on the event occurrences and co-occurrences in literature, without accounting for dynamic behavior, and therefore, ACCORDION outperforms it, as it is guided by the desired system behavior (i.e., the set of properties 𝒯 and their corresponding goal property probabilities 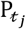).

## 4 Conclusions

In this paper, we have described a novel methodology and a tool, ACCORDION, that can be used to automatically assemble the information extracted from literature into models and to recommend models that achieve desired dynamic behavior. Our proposed approach combines machine reading with clustering, simulation, and model checking, into an automated framework for rapid model assembly and testing to address biological questions. Furthermore, by automatically extending models with the information published in literature, our methodology allows for efficient collection of the existing information in a consistent and comprehensive way, while also facilitating information reuse and data reproducibility, and often helping replace tedious trial-and-error manual experimentation, thereby increasing the pace of knowledge advancement. The results we presented here demonstrate different research scenarios where ACCORDION can be used. Both the benchmark set we presented, and the ACCORDION tool with detailed documentation are prepared for open access. As our next steps, we are planning to improve the input pre-processing in order to provide more useful candidate event sets, to make ACCORDION compatible with other model representation formats (e.g., SBML), as well as to work on parallelizing the tool implementation to improve the runtime when testing large number of properties.

## Acknowledgment

We would like to thank Kara Bocan, Khaled Sayed and Adam Butchy for useful discussions in early stages of the project.

