## Appendix for "Context-aware knowledge selection and reliable model recommendation with ACCORDION"

### Links to ACCORDION documentation and Jupyter notebook

https://accordion.readthedocs.io

https://mybinder.org/v2/gh/pitt-miskov-zivanov-lab/ACCORDION/HEAD?labpath=%2Fexamples%2Fuse_ACCORDION.ipynb

### ACCORDION notation and equations

In this section, we list the terms that were introduced in the main text, as well as the definitions of all the metrics used to evaluate the performance of ACCORDION.

*Baseline model* (BM) – The starting, initial model that is to be extended. The baseline model can be created manually with expert input, automatically from data, or adopted from models published in literature and in model databases.

*Golden model* (GM) – The "ground truth" model, manually created and rigorously curated.

*Candidate event* (CE) – Directed interactions between two elements (e.g., A and B), in the form "A influences B", where influences can be different biological mechanisms and can have positive or negative sign. CEs are used to extend the baseline model and they can be collected from different knowledge sources such as expert knowledge, published literature and pathway databases.

*Candidate model* (CM) – The model that ACCORDION assembles from the baseline model and a CE set.

*Node overlap* ($NO$) – The ratio of the number of nodes within a given cluster that are present in the baseline model and the total number of nodes within the cluster.

$\mathcal{T}$ – set of all properties

*Property* $t_j \in \mathcal{T}$ – a formal expression of actual, observed, measured, or expected behavior of the system.

*Property probability estimate* – $p_{t_j}^{\mathcal{M}}$ — a probability that model $\mathcal{M}$ satisfies property $t_j$.

*Goal property probability* – $\mathrm{P}_{t_j}$ –estimated or expected likelihood for the real system to satisfy property $t_j \in \mathcal{T}$

*Model property error* – $\varepsilon_{t_j}^{\mathcal{M}} = \left| p_{t_j}^{\mathcal{M}} - \mathrm{P}_{t_j} \right|$ – absolute error for model $\mathcal{M}$ in property $t_j$ as a difference between property probability estimate and goal property probability

*Average model error* – $\varepsilon_{\mathcal{T},avg}^{\mathcal{M}} = \frac{1}{|\mathcal{T}|} \sum_{t_j \in \mathcal{T}} \varepsilon_{t_j}^{\mathcal{M}}$ – computed for $\mathcal{M}$ across all properties $t_j \in \mathcal{T}$

*σ-score* – $\sigma_{\mathcal{T}}^{\mathcal{M}} = 1 - \varepsilon_{\mathcal{T},avg}^{\mathcal{M}}$ – computed for $\mathcal{M}$ with respect to a set of properties $\mathcal{T}$

*δ-limit* for $\varepsilon_{t_j}^{\mathcal{M}}$, that is, if $\varepsilon_{t_j}^{\mathcal{M}} \leq \delta$, then it is considered that $\mathcal{M}$ satisfies property $t_j$; (δ can have different values)

$\mathcal{D} = \left\{ t_j \mid t_j \in \mathcal{T} \text{ and } \varepsilon_{t_j}^{\mathcal{M}} \leq \delta \right\}$ – set of δ-satisfied properties; $\mathcal{D}$ is a subset of $\mathcal{T}$

*Absolute δ-score* – $N_{\mathcal{T},\delta}^{\mathcal{M}} = |\mathcal{D}|$ – number of δ-satisfied properties

*δ-score* – $N_{\mathcal{T},\delta,\%}^{\mathcal{M}} = \frac{N_{\mathcal{T},\delta}^{\mathcal{M}} * 100}{|\mathcal{T}|}$ – percent of properties within $\mathcal{T}$ that model $\mathcal{M}$ satisfies, assuming δ;

$\mathfrak{C}$ - set of all generated CMs

$\tilde{\sigma}_{\mathcal{T}}^{CM_l} = \left( \sigma_{\mathcal{T}}^{CM_l} - \min_{CM_i \in \mathfrak{C}} \left( \sigma_{\mathcal{T}}^{CM_i} \right) \right) / \left( \max_{CM_i \in \mathfrak{C}} \left( \sigma_{\mathcal{T}}^{CM_i} \right) - \min_{CM_i \in \mathfrak{C}} \left( \sigma_{\mathcal{T}}^{CM_i} \right) \right)$ – normalized $\sigma$-score

$\tilde{\varepsilon}_{\mathcal{T},avg}^{CM_l} = \left( \varepsilon_{\mathcal{T},avg}^{CM_l} - \min_{CM_i \in \mathfrak{C}} \left( \varepsilon_{\mathcal{T},avg}^{CM_i} \right) \right) / \left( \max_{CM_i \in \mathfrak{C}} \left( \varepsilon_{\mathcal{T},avg}^{CM_i} \right) - \min_{CM_i \in \mathfrak{C}} \left( \varepsilon_{\mathcal{T},avg}^{CM_i} \right) \right)$ – normalized average error

$\tilde{N}_{\mathcal{T},\delta,\%}^{CM_l} = \left( N_{\mathcal{T},\delta,\%}^{CM_l} - \min_{CM_i \in \mathfrak{C}} \left( N_{\mathcal{T},\delta,\%}^{CM_i} \right) \right) / \left( \max_{CM_i \in \mathfrak{C}} \left( N_{\mathcal{T},\delta,\%}^{CM_i} \right) - \min_{CM_i \in \mathfrak{C}} \left( N_{\mathcal{T},\delta,\%}^{CM_i} \right) \right)$ – normalized δ-score

$\tilde{p}_{\mathcal{T}}^{CM_l} = \left(\prod_{t_j \in \mathcal{T}} p_{t_j}^{CM_l} - \min_{CM_i \in \mathfrak{C}} \left(\prod_{t_j \in \mathcal{T}} p_{t_j}^{CM_i}\right)\right) / \left(\max_{CM_i \in \mathfrak{C}} \left(\prod_{t_j \in \mathcal{T}} p_{t_j}^{CM_i}\right) - \min_{CM_i \in \mathfrak{C}} \left(\prod_{t_j \in \mathcal{T}} p_{t_j}^{CM_i}\right)\right)$ – normalized joint probability of satisfying all properties in $\mathcal{T}$ (assuming independence in properties).

### Case studies

We describe here the creation of nine case studies, three studies for each system, naïve T cell differentiation, T cell large granular lymphocyte leukemia, and pancreatic cancer cells. In particular, we provide the details of baseline models, golden models, candidate event (CE) sets, properties and scenarios.

For all the nine case studies, we used the PubMed database [1]. The PubMed search was conducted using Entrez [2], an integrated database retrieval system that allows access to a diverse set of databases at the National Center for Biotechnology Information (NCBI) website. The published articles that were obtained through search of PubMed are read using the REACH engine [3], which extracted a list of events and the corresponding information. The REACH reading engine is available online and can be run through the Integrated Network and Dynamical Reasoning Assembler (INDRA) [4].

#### *T cell differentiation (Tcell)*

**Background**: Naïve peripheral T cells are stimulated via antigen presentation to T cell receptor (TCR) and with co-stimulation at CD28 receptor. This stimulation results in the activation of several downstream pathways, and feedback and feedforward loops between pathway elements, which then lead to the differentiation of naïve T cells into helper (Th) or regulatory (Treg) phenotypes. The distribution between Th and Treg cells within the T cell population depends on antigen dose; for instance, high antigen dose results in prevalence of Th cells, while low antigen dose leads to a mixed population of Th and Treg cells. The key markers that are commonly used to measure the outcomes of the naïve T cell differentiation into Th and Treg cells are IL2 and Foxp3, respectively. In other words, Th cells are characterized by high expression of IL-2 and low expression of Foxp3, and Treg cells are characterized by high expression of Foxp3 and low expression of IL-2.

**Baseline and Golden models**: In [5], the authors proposed a model where most of the elements are assumed to have two main levels of activity, and are therefore represented with Boolean variables, and their update rules are logic functions. Additionally, the stimulation through TCR is assumed to have three different levels, no stimulation (TCR=0), low dose (TCR=1), and high dose (TCR=2), and therefore, it is implemented using two Boolean variables. In [6], the authors have proposed an extension of the original T cell model from [5], a new model that improved the behavior of the original model. Specifically, in the new model in [6], the Foxp3 response to low dose is closer to experimental observations, that is, it is present in almost 70% of the differentiated population, while in [5] Foxp3 was present in 100% of the differentiated population. In both models, there is a brief transient induction of Foxp3 after the stimulation with high antigen dose. We refer to the model from [6] as the golden model. As the baseline model, we used the original model from [5], without several interactions overlapping with the golden model from [6] (TCR activates PIP3, PIP3 activates Akt, Akt activates mTORC2 and mTORC2 inhibits Akt).

**Properties**: From the golden model in [6] and the results of its studies, we define a set of properties that the final model recommended by ACCORDION should satisfy. Specifically, the properties capture observed responses of key pathway components in T cells, Foxp3, IL-2, PTEN, CD25, STAT5, AKT, mTOR, mTORC2 and FoxO1, to three scenarios: (1) no stimulation (TCR=0), (2) stimulation with low antigen dose (TCR=1), and (3) stimulations with high antigen dose (TCR=2). The complete list of 27 properties is shown in Table S1. While the model from [5] satisfied a large number of system properties, except for a few that are satisfied by the model in [6] only, the baseline model in its reduced shape does not satisfy a larger set of system properties.

**Candidate event sets**: The CE set, which is another input to ACCORDION, is assembled in three different ways for the T cell case studies. In the *Fully Automated CE set* (Tcell $CE^{FA}$), both the PubMed database search for relevant articles and the extraction of event data from the selected articles were done by machines. In this first experiment, we used the search query "*T-cell and (PTEN or AKT or FOXO)*" and selected top 11 from the best matched papers, by the PubMed search engine. In the *Semi-Automated CE set* (Tcell $CE^{SA}$), we selected papers that are cited by [6] and used the event information that REACH automatically extracted from those papers. Finally, in the *Semi-Mannual CE set* (Tcell $CE^{SM}$), we excluded from the $CE^{SA}$ set those interactions that violate any assumptions made by the authors originally in [5]. For instance, the authors in [5] consider element TCR to be an input to the network, and therefore, TCR should not have any regulators in the T cell model. Therefore, if REACH retrieves an interaction in which TCR is a regulated element, we remove these interactions and keep only the interactions having TCR as a regulator. We have also selected from the reading output the protein-protein interactions, not including the more general biological processes information. The rationale behind this is that there is often not enough context for a mentioned biological process, and the lack of context affects interpretation of the extracted interaction. The machine reading outputs obtained by reading papers for each case contain 239, 131 and 85 interactions for the Tcell $CE^{FA}$, Tcell $CE^{SA}$ and Tcell $CE^{SM}$ cases, respectively.

#### *T cell large granular lymphocyte (T-LGL) leukemia*

**Background**: The blood cancer T cell large granular lymphocyte (T-LGL) leukemia is a chronic disease characterized by an abnormal increase of T cells [7]. There is no curative therapy yet known for this disease. Hence, there is a crucial need to identify potential therapeutic targets. A discrete dynamic model of the disease was published in [7]. Eventually, the authors in [8] performed a comprehensive dynamical and structural analysis of the network model which led to the identification of 19 therapeutic targets [8].

**Baseline and Golden models**: For this case study, the model published in [7] serves as the golden model, whereas the baseline model is created by removing all direct regulators of the 19 model elements that were identified by [8] as therapeutic targets. According to [8], these elements are: BID, Caspase, Ceramide, DISC, ERK, GAP, IL2RB, IL2RBT, JAK, MCL1, MEK, NFKB, PDGFR, PI3K, RAS, S1P, SOCS, SPHK1 and STAT3.

**Properties**: The properties that our final automatically extended model needs to satisfy capture the observed responses of the 19 therapeutic targets in the golden model under one scenario (Table S1).

**Queries and Candidate event sets**: Next, we came up with three different queries that are the input to the search engine in order to retrieve the most relevant sets of papers. The first T-LGL case study used simple query "*T-LGL leukemia therapeutic targets and apoptosis*" (T-LGL $Q^{Sm}$). From the papers that PubMed returned, we then selected 22 papers that PubMed identified as "Best match". The machine reading output obtained by reading those papers contains 52 interactions. The second and third T-LGL case studies used medium query "*T cell large granular lymphocyte (T-LGL) leukemia proliferation apoptosis*" (T-LGL $Q^{Med}$) and detailed query "*T cell large granular lymphocyte (T-LGL) leukemia therapeutic targets proliferation apoptosis*" (T-LGL $Q^{Det}$). The number of interactions used in these case studies are 448 and 644 extracted from 38 and 46 papers, respectively. As can be noticed, the queries were designed so that they include key words related to the T-LGL model and therapeutic targets. For each query and the corresponding set of papers, we used REACH to read these papers and extract the set of CEs as input to ACCORDION.

#### *Pancreatic cancer cell (PCC)*

**Background**: The modeling of pancreatic cancer is of great importance since it could reveal some molecular mechanisms that help in guiding treatment. In [9], the authors created manually a discrete model of the major signaling pathways, metabolism and the tumor microenvironment including macrophages. The model is initialized with pancreatic cancer receptors and mutations and simulated in time. The model describes the hallmarks of cancer and suggests combinations of inhibitors as therapies. These hallmarks are represented as the processes of apoptosis, autophagy, cell cycle progression, inflammation, immune response, oxidative phosphorylation and proliferation.

**Baseline and Golden models**: The model in [9] serves as a golden model for our PCC case studies. We removed from the model in [9] a subset of paths that have autophagy, apoptosis and proliferation as their target nodes to create three different baseline models for the PCC case studies. Unlike the T-LGL case study where we used the same baseline model and different CEs, here, we designed each experiment with a different baseline model. This is achieved by removing the paths that connect a source node that initiates a specific biological process such as (autophagy, apoptosis and proliferation) to a target node which is this biological process. For instance, according to the evidence that mTORC1 initiates autophagy [10], we remove the paths that link mTORC1 and autophagy and the outcome is our first baseline model and a corresponding PCC case study (PCC $BM^{Au}$). Similarly, based on the fact that TGFβ1 regulates apoptosis [11] and KRas mutations enhances proliferation [12], we have also created two additional baseline models and corresponding PCC case studies by removing the paths that connect source nodes to target nodes, TGFβ1 to apoptosis (PCC $BM^{Ap}$ case study) and KRas to proliferation (PCC $BM^{Pr}$ case study), respectively. In all three PCC case studies, the golden model is the whole PCC model network [9].

**Properties**: Using the golden model and the descriptions from [9], we wrote the BLTL expressions of 21 system properties that capture the behavior of seven key elements (apoptosis, autophagy, cell cycle progression, immune response, inflammation, oxidative phosphorylation and proliferation) under three different scenarios, (1) normal, (2) with injury and (3) with KRas, TP53 and CDN2A mutation [13] (Table S1).

**Candidate event sets**: The CE input set for ACCORDION is the same for the three PCC case studies, it contains 631 interactions retrieved from 19 papers cited in the PCC model paper [9].

*Table S1* The set of properties that are observed to be true in the T cells, T-LGL cells and pancreatic cancer cells, for several scenarios, written in plain English and in BLTL format, and their goal probability values ($P_{t_j}$).

| Scenario | Prop. # | Description | BLTL format | Expected value |
|---|---|---|---|---|
| No TCR | 1 | Once deactivated, AKT will remain inactive until end of analyzed period | F[1500]G[50](AKT==0) | 1 |
| | 2 | Once activated, PTEN will remain active until end of analyzed period | F[1500]G[50](PTEN==1) | 0.338 |
| | 3 | Once deactivated, FOXP3 will remain inactive until end of analyzed period | F[1500]G[50](FOXP3==0) | 1 |
| | 4 | Once deactivated, IL2 will remain inactive until end of analyzed period | F[1500]G[50](IL2==0) | 1 |
| | 5 | Once deactivated, CD25 will remain inactive until end of analyzed period | F[1500]G[50](CD25==0) | 1 |
| | 6 | Once deactivated, STAT5 will remain inactive until end of analyzed period | F[1500]G[50](STAT5==0) | 1 |
| | 7 | Once deactivated, mTORC1 will remain inactive until end of analyzed period | F[1500]G[50](MTORC1==0) | 1 |
| | 8 | Once deactivated, mTORC2 will remain inactive until end of analyzed period | F[1500]G[50](MTORC2==0) | 1 |
| | 9 | Once activated, FOXO1 will remain active until end of analyzed period | F[1500]G[50](FOXO1==1) | 1 |
| Low TCR | 10 | Once deactivated, AKT will remain inactive until end of analyzed period | F[1500]G[50](AKT==0) | 1 |
| | 11 | Once activated, PTEN will remain active until end of analyzed period | F[1500]G[50](PTEN==1) | 0.9 |
| | 12 | Once activated, FOXP3 will remain active until end of analyzed period | F[1500]G[50](FOXP3==1) | 0.760 |
| | 13 | Once deactivated, IL2 will remain inactive until end of analyzed period | F[1500]G[50](IL2==0) | 1 |
| | 14 | Once activated, CD25 will remain active until end of analyzed period | F[1500]G[50](CD25==1) | 1 |
| | 15 | Once activated, STAT5 will remain active until end of analyzed period | F[1500]G[50](STAT5==1) | 1 |
| | 16 | Once deactivated, mTORC1 will remain active until end of analyzed period | F[1500]G[50](MTORC1==0) | 1 |
| | 17 | Once activated, mTORC2 will remain active until end of analyzed period | F[1500]G[50](MTORC2==1) | 1 |
| | 18 | Once activated, FOXO1 will remain active until end of analyzed period | F[1500]G[50](FOXO1==1) | 1 |
| High TCR | 19 | Once activated, AKT will remain inactive until end of analyzed period | F[1500]G[50](AKT==1) | 1 |
| | 20 | In developing Th, PTEN decreases and remains absent | F[1500]G[50](PTEN==0) | 1 |
| | 21 | Once deactivated, FOXP3 will remain inactive until end of analyzed period | F[1500]G[50](FOXP3==0) | 1 |
| | 22 | Once activated, IL2 will remain active until end of analyzed period | F[1500]G[50](IL2==1) | 1 |
| | 23 | Once activated, CD25 will remain active until end of analyzed period | F[1500]G[50](CD25==1) | 1 |
| | 24 | Once activated, STAT5 will remain active until end of analyzed period | F[1500]G[50](STAT5==1) | 1 |
| | 25 | Once activated, mTORC1 will remain inactive until end of analyzed period | F[1500]G[50](MTORC1==1) | 1 |
| | 26 | Once activated, mTORC2 will remain active until end of analyzed period | F[1500]G[50](MTORC2==1) | 1 |
| | 27 | Once activated, FOXO1 will remain active until end of analyzed period | F[1500]G[50](FOXO1==0) | 1 |

| Scenario | Prop.# | Description | BLTL format | Expected value |
| --- | --- | --- | --- | --- |
| Normal | 1 | Once deactivated, apoptosis will remain inactive until end of analyzed period | F[9500]G[50](apoptosis==0) | 1 |
| | 2 | Once deactivated, autophagy will remain inactive until end of analyzed period | F[9500]G[50](autophagy==0) | 1 |
| | 3 | Once deactivated, cell cycle progression will remain inactive until end of analyzed period | F[9500]G[50](cell cycle progression==0) | 1 |
| | 4 | Once deactivated, immune response will remain inactive until end of analyzed period | F[9500]G[50](immune response==0) | 1 |
| | 5 | Once deactivated, inflammation will remain inactive until end of analyzed period | F[9500]G[50](inflammation==0) | 1 |
| | 6 | Once deactivated, oxidative phosphorylation will remain inactive until end of analyzed period | F[9500]G[50](oxidative phosphorylation==0) | 1 |
| | 7 | Once deactivated, proliferation will remain inactive until end of analyzed period | F[9500]G[50](proliferation==0) | 1 |
| With injury | 9 | Once deactivated, apoptosis will remain inactive until end of analyzed period | F[9500]G[50](apoptosis==0) | 1 |
| | 9 | Once deactivated, autophagy will remain inactive until end of analyzed period | F[9500]G[50](autophagy==0) | 1 |
| | 10 | Once deactivated, cell cycle progression will remain inactive until end of analyzed period | F[9500]G[50](cell cycle progression==0) | 1 |
| | 11 | Once deactivated, immune response will remain inactive until end of analyzed period | F[9500]G[50](immune response==0) | 1 |
| | 12 | Once activated, inflammation will remain active until end of analyzed period | F[9500]G[50](inflammation==1) | 1 |
| | 13 | Once activated, oxidative phosphorylation will remain active until end of analyzed period | F[9500]G[50](oxidative phosphorylation==1) | 1 |
| | 14 | Once deactivated, proliferation will remain inactive until end of analyzed period | F[9500]G[50](proliferation==0) | 1 |
| With KRas, TP53, CDN2A mutation | 15 | Once deactivated, apoptosis will remain inactive until end of analyzed period | F[9500]G[50](apoptosis==0) | 1 |
| | 16 | Once activated, autophagy will remain active until end of analyzed period | F[9500]G[50](autophagy==1) | 1 |
| | 17 | Once activated, cell cycle progression will remain active until end of analyzed period | F[9500]G[50](cell cycle progression==1) | 1 |
| | 18 | Once deactivated, immune response will remain inactive until end of analyzed period | F[9500]G[50](immune response==0) | 1 |
| | 19 | Once activated, inflammation will remain active until end of analyzed period | F[9500]G[50](inflammation==1) | 1 |
| | 20 | Once deactivated, oxidative phosphorylation will remain inactive until end of analyzed period | F[9500]G[50](oxidative phosphorylation==0) | 1 |
| | 21 | Once activated, proliferation will remain active until end of analyzed period | F[9500]G[50](proliferation==1) | 1 |

| Scenario | Prop. # | Description | BLTL format | Expected value |
| --- | --- | --- | --- | --- |
| | 1 | Once deactivated, DISC will remain inactive until end of analyzed period | F[1500]G[50](DISC==0) | 1 |
| | 2 | Once deactivated, Ceramide will remain inactive until end of analyzed period | F[1500]G[50](Ceramide==0) | 1 |
| | 3 | Once deactivated, Caspase will remain inactive until end of analyzed period | F[1500]G[50](Caspase==0) | 1 |
| | 4 | Once activated, SPHK1 will remain active until end of analyzed period | F[1500]G[50](SPHK1==1) | 1 |
| | 5 | Once activated, S1P will remain active until end of analyzed period | F[1500]G[50](S1P==1) | 1 |
| | 6 | Once activated, PDGFR will remain active until end of analyzed period | F[1500]G[50](PDGFR==1) | 1 |
| | 7 | Once deactivated, GAP will remain inactive until end of analyzed period | F[1500]G[50](GAP==0) | 1 |
| | 8 | Once activated, RAS will remain active until end of analyzed period | F[1500]G[50](RAS==1) | 1 |
| | 9 | Once activated, MEK will remain active until end of analyzed period | F[1500]G[50](MEK==1) | 1 |
| | 10 | Once activated, ERK will remain active until end of analyzed period | F[1500]G[50](ERK==1) | 1 |
| | 11 | Once activated, IL2RBT will remain active until end of analyzed period | F[1500]G[50](IL2RBT==1) | 1 |
| | 12 | Once activated, IL2RB will remain active until end of analyzed period | F[1500]G[50](IL2RB==1) | 1 |
| | 13 | Once activated, STAT3 will remain active until end of analyzed period | F[1500]G[50](STAT3==1) | 1 |
| | 14 | Once deactivated, BID will remain inactive until end of analyzed period | F[1500]G[50](BID==0) | 1 |
| | 15 | Once activated, MCL1 will remain active until end of analyzed period | F[1500]G[50](MCL1==1) | 1 |
| | 16 | Once deactivated, SOCS will remain inactive until end of analyzed period | F[1500]G[50](SOCS==0) | 1 |
| | 17 | Once activated, JAK will remain active until end of analyzed period | F[1500]G[50](JAK==1) | 1 |
| | 18 | Once activated, PI3K will remain active until end of analyzed period | F[1500]G[50](PI3K==1) | 1 |
| | 19 | Once activated, NFKB will remain active until end of analyzed period | F[1500]G[50](NFKB==1) | 1 |
